# First molecular evidence that perialgal vacuole membrane maturation is temporally regulated during establishment of *Chlorella variabilis* photoendosymbiosis in *Paramecium tritobursaria*

**DOI:** 10.64898/2026.08.14.744893

**Authors:** Yuuki Kodama, Masahiro Fujishima

## Abstract

Photoendosymbiosis between the ciliate *Paramecium tritobursaria* and the green alga *Chlorella variabilis* provides a model for understanding stable photoendosymbiosis. A defining feature of this association is the perialgal vacuole (PV) membrane, a host-derived membrane that encloses each alga and prevents its digestion. However, the timing of PV membrane maturation remains poorly understood because of the lack of molecular markers to distinguish between immature and mature PV membranes. Previous studies have shown that the establishment of symbiosis proceeds through multiple regulated steps following algal uptake; however, the molecular maturation of the PV membrane has not been directly examined. Here, we report a monoclonal antibody that specifically recognizes the PV membrane in symbiotic *P. tritobursaria*. Time-course immunofluorescence analysis showed that the PV membrane antigen was absent in the early stages after algal uptake, appeared at 48 h, and was detected in all PV membranes by 72 h. The antigen persisted before and after synchronous PV swelling, an experimentally inducible state associated with the loss of normal PV membrane function, but was absent from the membranes surrounding the digested algae. Our findings provide the first molecular evidence that PV membrane maturation is a temporally regulated checkpoint during the establishment of photoendosymbiosis.

## 1 Introduction

The ciliate *Paramecium tritobursaria* (*P. tritobursaria*) is an established experimental model for photoendosymbiosis, as it forms a facultative and readily re establishable mutualism with endosymbiotic green algae, allowing both partners to live independently (Siegel and Karakashian, 1959; Karakashian and Karakashian, 1965; Reisser, 1987; Kodama et al., 2014; Song et al., 2017; He et al., 2019; Cheng et al., 2020; Jenkins, 2024). In this system, a perialgal vacuole (PV) derived from the host digestive vacuole (DV) membrane plays a central role in symbiont protection and metabolic interactions between the host and algae (Karakashian et al., 1968; Reisser, 1986). The defining structural features of the PV membrane are that it individually encloses each alga and is stably positioned immediately beneath the host cell cortex (Reisser, 1992). The PV membrane is a compartment distinct from the DV, as demonstrated by freeze-fracture and ultrastructural studies, which have revealed clear morphological differences, including a consistent structure and high intramembranous particle density, distinguishing PV membranes from DV membranes (Meier et al., 1980; Meier et al., 1984). These observations establish the PV membrane as a specialized symbiotic interface, rather than a modified lysosomal membrane. Our current knowledge of indicates that microalgae are commonly engulfed via phagocytosis and subsequently retained intracellularly within a host-derived vacuole, termed the symbiosome, which is widely regarded as an arrested phagosome (Davy et al., 2012). Functionally, a hallmark of the mature PV membrane is its resistance to lysosomal fusion, thereby preventing the digestion of enclosed symbionts (Karakashian and Karakashian, 1973; Karakashian and Rudzinska, 1981). Comparative studies across diverse photosymbiotic systems have further shown that healthy algal symbionts evade host digestive pathways after phagocytosis, whereas damaged or non-viable symbionts are targeted for lysosomal degradation (Davy et al., 2012). Environmental conditions strongly influence the fate of symbiotic algae; under light conditions, *Chlorella* spp. symbionts remain intact and contribute to host metabolism, whereas under prolonged darkness, they gradually degenerate and are ultimately digested (Gu et al., 2002). Enzyme cytochemistry revealed elevated activities of acid phosphatase (AcPase) and other hydrolases in dark-cultured *P. bursaria*, linking enhanced digestive activity to symbiont breakdown (Gu et al., 2002). In addition, the PV lumen is maintained under acidic conditions (Schüßler and Schnepf, 1992), which promotes light-dependent maltose release from symbiotic algae into the host cell (Shibata et al., 2016). Together, these findings highlight that the PV membrane is a critical interface for integrating symbiont protection and metabolic exchange. Recent comparative reviews have further emphasized that such symbiotic compartments create a highly controlled intracellular microenvironment that supports algal metabolism and long-term persistence across diverse freshwater and marine photoendosymbioses (Yee et al., 2025). Accumulating evidence suggests that the establishment and maintenance of photoendosymbiosis in *P. bursaria* is accompanied by marked changes in mitochondrial abundance, activity, and spatial organization, including their positioning around symbiotic algae, indicating close functional coordination between host mitochondria and the PV membrane (Fujishima and Kodama, 2012; Song et al., 2017; Kodama and Fujishima, 2022; Fujishima and Nishiyama, 2026; Hayakawa and Suzaki, 2026). These observations raise the possibility that the acquisition of a mature PV membrane is temporally coordinated with the metabolic remodeling of the host cell.

Despite more than half a century of intensive cytological, physiological, and biochemical research, the molecular basis of PV membrane formation and maturation remains unclear. One major obstacle is the absence of molecular markers applicable to highly divergent, non-model protists, for which commercially available antibodies provide little specificity. Previous biochemical studies have identified PV-associated glycoproteins, including Concanavalin A-binding proteins (Hiraoka et al., 2008), and recent high-resolution chemical imaging using OrbiSIMS has revealed PV-specific oligosaccharide and lipid signatures (Aoyagi et al., 2019). Furthermore, the isolation of symbiotic algae retaining intact PV membranes has been achieved, with the presence of the PV membrane was confirmed by transmission electron microscopy (TEM) (Omura et al., 2007). More recently, (Fujishima and Nishiyama, 2026) succeeded in isolating PV membranes as part of the mitochondria– PV membrane complexes and directly visualized these complexes using lipophilic fluorescent probes, providing clear structural evidence for the physical integrity and spatial organization of the PV membrane. Although these studies provide strong structural evidence that the PV membrane represents a distinct host-derived compartment with a unique molecular identity, they leave unresolved the central question: when and how does the PV membrane acquire its mature symbiotic properties during the establishment of symbiosis?

Extensive analyses of artificial re-photoendosymbiosis in alga-free *P. bursaria* have demonstrated that stable endosymbiosis is not an automatic consequence of algal uptake but proceeds through a series of tightly regulated cellular processes (Kodama and Fujishima, 2008b; 2010; Kodama and Fujishima, 2024; Fujishima and Nishiyama, 2026). Based on detailed cytological and ultrastructural observations, this process can be conceptually divided into four sequential phenomena (Kodama and Fujishima, 2010; Kodama and Fujishima, 2024): (i) temporary resistance of internalized algae to host digestive enzymes within DVs, (ii) escape of individual algae from DVs into the host cytoplasm via DV membrane budding, (iii) transformation of the DV membrane into a PV membrane that encloses each alga individually, and (iv) stable positioning of the PV membrane beneath the host cell cortex, where host mitochondria and trichocysts are localized. Although processes (i) and (ii), as well as the cellular consequences of process (iv), have been described in considerable detail, the molecular maturation of the PV membrane during process (iii), particularly the timing at which it acquires its definitive symbiotic identity, has never been directly examined.

Previous studies have demonstrated that PV membranes rapidly lose lysosomal characteristics shortly after algal escape from DVs during the early post-uptake stage, preceding stable infection (Kodama and Fujishima, 2009). However, whether and when the PV membrane subsequently acquires its mature molecular identity remains unclear.

In this study, we report the first monoclonal antibody (mAb) that specifically recognizes the PV membrane in *P. tritobursaria*. Using this molecular marker, we examined PV membrane behavior during symbiosis establishment and during synchronous PV swelling (SPVS), an experimentally inducible state that reflects the loss of normal PV membrane function (Kodama and Fujishima, 2008a). Specifically, we (i) optimized fixation methods for indirect immunofluorescence microscopy, (ii) demonstrated antigen specificity by comparing algae-bearing and alga-free *P. tritobursaria* cells, (iii) determined the time-dependent appearance of the PV membrane antigen during experimental infection, and (iv) examined changes in antigen retention before and after SPVS, including the membranes surrounding the digested algae. Using this approach, we addressed whether PV membrane maturation is a temporally regulated process following algal uptake, providing new insights into the molecular differentiation of the symbiotic membranes.

## 2 Materials and methods

### 2.1 Cultivation of alga-free and algae-bearing *P. tritobursaria*

The cells used in this study were identified as *P. tritobursaria* according to the classification of Spanner et al. (2022) (Spanner et al., 2022), corresponding to *P. bursaria* syngen 1 in the classification of Bomford (1966) (Bomford, 1966). Two *P. tritobursaria* strains were used in this study: the alga-free (aposymbiotic) strain Yad1w and the algae-bearing strain Yad1g1N (mating type I). The Yad1g1N strain was originally established by infecting alga-free Yad1w cells with the symbiotic *C. variabili*s strain 1N (Kodama and Fujishima, 2011) and has been continuously maintained under laboratory culture conditions for more than 14 years. Therefore, Yad1g1N cells represent a long-term, fully stabilized model of mature photoendosymbiosis rather than a recently infected or transitional state.

*P. tritobursaria* cells were cultured in red pea (*Pisum sativum*) extract medium (Tsukii et al., 1995) prepared in modified Dryl’s solution (MDS), in which NaH_2_PO_4_·2H_2_O was replaced with KH_2_PO_4_ (Dryl, 1959). The bacterium *Klebsiella aerogenes* (ATCC 35028) was added as a food source 1 day before use to red pea extract medium (Fujishima et al., 1990). Several hundred *P. tritobursaria* cells were used to inoculate 2 mL aliquots of culture medium in test tubes, followed by the daily addition of 2 mL of fresh medium for 12 days. Cultures were maintained at 23 ± 1 °C and used for experiments 1 day after the final feeding, corresponding to the early stationary phase of growth. Algae-bearing Yad1g1N cells were cultured under constant illumination (24L:0D) using fluorescent light (27 W, natural white) at an intensity of 20–30 μmol photons m ² s ¹.

### 2.2 Production of monoclonal antibody

A monoclonal antibody (mAb), clone 1H2E11G7H1, was generated and used in this study. Mass cultures of algae-bearing *P. tritobursaria* cells in the early stationary phase of growth were harvested and homogenized. The homogenate was intraperitoneally injected into 8-week-old BALB/c mice. Hybridoma cell lines were generated using the method described by (Galfrè and Milstein, 1981) and (Fujishima et al., 1992). Culture supernatants were screened using indirect immunofluorescence microscopy, and clones producing antibodies specific to the PV membrane were selected. Eight-week-old BALB/c mice used in this study were obtained from CLEA Japan, Inc. (Tokyo, Japan). Hybridoma production for monoclonal antibody generation was initiated before the establishment of the current university-wide animal ethics committee system at Yamaguchi University; therefore, no institutional approval number is available. All animal procedures were performed in accordance with accepted ethical standards and established practices for the care and use of laboratory animals in Japan at the time the experiments were conducted and were supervised by the corresponding author. All efforts were made to minimize animal suffering and to use the minimum number of animals required for antibody production.

### 2.3 Indirect immunofluorescence microscopy

Several hundred *P. tritobursaria* cells were air-dried on coverslips (4.5 × 24 mm; Matsunami Glass, Ltd., Osaka, Japan) and fixed for 10 min at 4 °C with either 4% (w/v) paraformaldehyde (PFA) in phosphate-buffered saline (PBS; 137 mM NaCl, 2.68 mM KCl, 8.1 mM Na_2_HPO_4_·12H_2_O, 1.47 mM KH_2_PO_4_, pH 7.2) or cold 99.5% (v/v) ethanol. After fixation, the cells were treated with PBST (PBS containing 0.05% (v/v) Tween-20) for 10 min at 4 °C. The cells were then washed twice with PBS for 10 min each at 4 °C and incubated with mAb (1:20 dilution) overnight at 4 °C. After primary antibody incubation, the cells were washed twice with PBS for 10 min each at room temperature and subsequently incubated with Alexa Fluor 488-conjugated goat anti-mouse IgG secondary antibody (Molecular Probes, 1:1,000 dilution) for 2 h at room temperature or overnight at 4 °C. After secondary antibody incubation, the cells were washed twice with PBS for 10 min each at room temperature. Samples were observed under a differential interference contrast (DIC) and fluorescence microscope (BX53; EVIDENT, Tokyo, Japan) equipped with U-FBW, U-FBNA, and U-FGW filter sets. The same procedure was applied to symbiotic *C. variabilis* cells isolated from the algae-bearing Yad1g1N strain. The red autofluorescence of *Chlorella* sp. cells was used as an intrinsic marker of symbiotic algae and merged with green immunofluorescence signals of the PV membrane antigen (Alexa Fluor 488) for image analysis.

### 2.4 Isolation of symbiotic *C. variabili*s from algae-bearing *P. tritobursaria*

Symbiotic *C. variabilis* strain 1N cells were isolated from algae-bearing *P. tritobursaria* Yad1g1N cultures (300 mL). The cultures were passed through two layers of Kimwipes (Kimberly-Clark Professional, Roswell, GA, USA) to remove debris and subsequently filtered through a 15-μm nylon mesh fitted to a centrifuge tube. The retained cells were washed with MDS, partially concentrated by filtration, and finally collected using a hand-operated centrifuge (UKG-2; Uchida Rikakiki, Tokyo, Japan), and resuspended in 1 mL of MDS. The suspension was supplemented with 0.1 mM phenylmethylsulfonyl fluoride (Sigma-Aldrich) and homogenized on ice with 30 strokes using a 1 mL-Teflon homogenizer on ice. The homogenate was filtered through a 15-μm nylon mesh, and the filtrate was washed three times with MDS by centrifugation at 4,350 × g for 1 min at room temperature. The final pellet was resuspended in 500 μL of MDS, and the algal cell density was determined using a Thoma hemocytometer.

### 2.5 Infection of alga-free *P. tritobursaria* cells with symbiotic *C. variabilis*

Isolated symbiotic *C. variabilis* 1N cells were mixed with alga-free *P. tritobursaria* at a ratio of 5 × 10 algal cells to 5 × 10³ host cells per mL in a 10 mL centrifuge tube. At 1.5 min after mixing, host cells were separated from uningested algae using a 15 μm nylon mesh and extensively washed with 100 mL of MDS to completely remove non ingested algae, followed by resuspension in MDS. The cells were maintained under constant light (24L:0D) at 20–30 μmol photons m ² s ¹ using fluorescent light (27 W, natural white). Aliquots (200 μL) were collected at 72 h to assess infection efficiency and fixed with an equal volume of 8% (w/v) PFA for DIC microscopy. For indirect immunofluorescence microscopy, aliquots collected at 10−30 min, 1.5−2.5, 24, 48, and 72 h were placed on coverslips (4.5 × 24 mm), air-dried, fixed with cold 99.5% (v/v) ethanol for 10 min, and processed for immunostaining. The cells were observed using DIC and fluorescence microscopy (BX53; EVIDENT), and images were captured using a DP74 camera. All fluorescence images were acquired with a fixed exposure time of 260 ms for quantitative comparison.

### 2.6 SPVS induction protocol

Synchronous PV swelling (SPVS) was induced as previously described by (Kodama and Fujishima, 2008a). Cycloheximide stock solutions (10 mg mL ¹ in distilled water; Wako Pure Chemical Industries, Ltd., Japan) were stored at −20 °C until use and diluted with MDS immediately before treatment to a final concentration of 10 μg mL ¹. Algae-bearing Yad1g1N cells were incubated with cycloheximide under constant light (24L:0D) at 20–30 μmol photons m ² s ¹ using fluorescent light (27 W, natural white). For indirect immunofluorescence microscopy, samples were collected 2 days after treatment and placed on coverslips (4.5 × 24 mm). The cells were air-dried and fixed with cold 99.5% (v/v) ethanol for 10 min. Fixed samples were processed for indirect immunofluorescence microscopy, as described above. In parallel experiments, a subset of cycloheximide-treated cells was observed without fixation using a DIC microscope, because SPVS cannot be visualized after fixation.

### 2.7 Statistical analysis

The proportion of PV membranes showing positive immunofluorescence was calculated for each time point. Exact 95% confidence intervals (CI) were calculated using the Clopper–Pearson exact method based on the binomial distribution. The kinetics of antigen appearance were visualized using graphs with confidence intervals.

## 3 Results

### 3.1 Effect of fixation methods on the visualization of the PV membrane antigen

To determine whether fixation methods influence the visualization of the PV membrane antigen, Yad1g1N cells were fixed with either 4% (w/v) paraformaldehyde (PFA) or 99.5% (v/v) ethanol (Figure 1). DIC images revealed no obvious morphological differences between the two fixation methods (Figures 1A, C). However, indirect immunofluorescence microscopy revealed clear differences in antigen visualization. In PFA-fixed cells (Figure 1B), the fluorescence signals appeared diffuse, and the PV membrane surrounding each alga could not be clearly resolved in the images. In contrast, ethanol-fixed cells exhibited sharply defined ring-like fluorescence around individual endosymbiotic algae (Figure 1D), allowing for the accurate visualization and enumeration of PV membranes. Despite these differences in clarity, both fixation methods indicated that the antigen was localized specifically to the PV membrane and not to other host structures. Because ethanol fixation provided superior resolution and consistency, all subsequent immunofluorescence experiments were performed using 99.5% ethanol. At higher magnification, ethanol-fixed cells showed continuous PV membrane antigen signals outlining individual symbiotic algae (Figures 1E–H). When the focal plane was adjusted to the algal surface, the PV membrane antigen occasionally appeared punctate.

**FIGURE 1.**
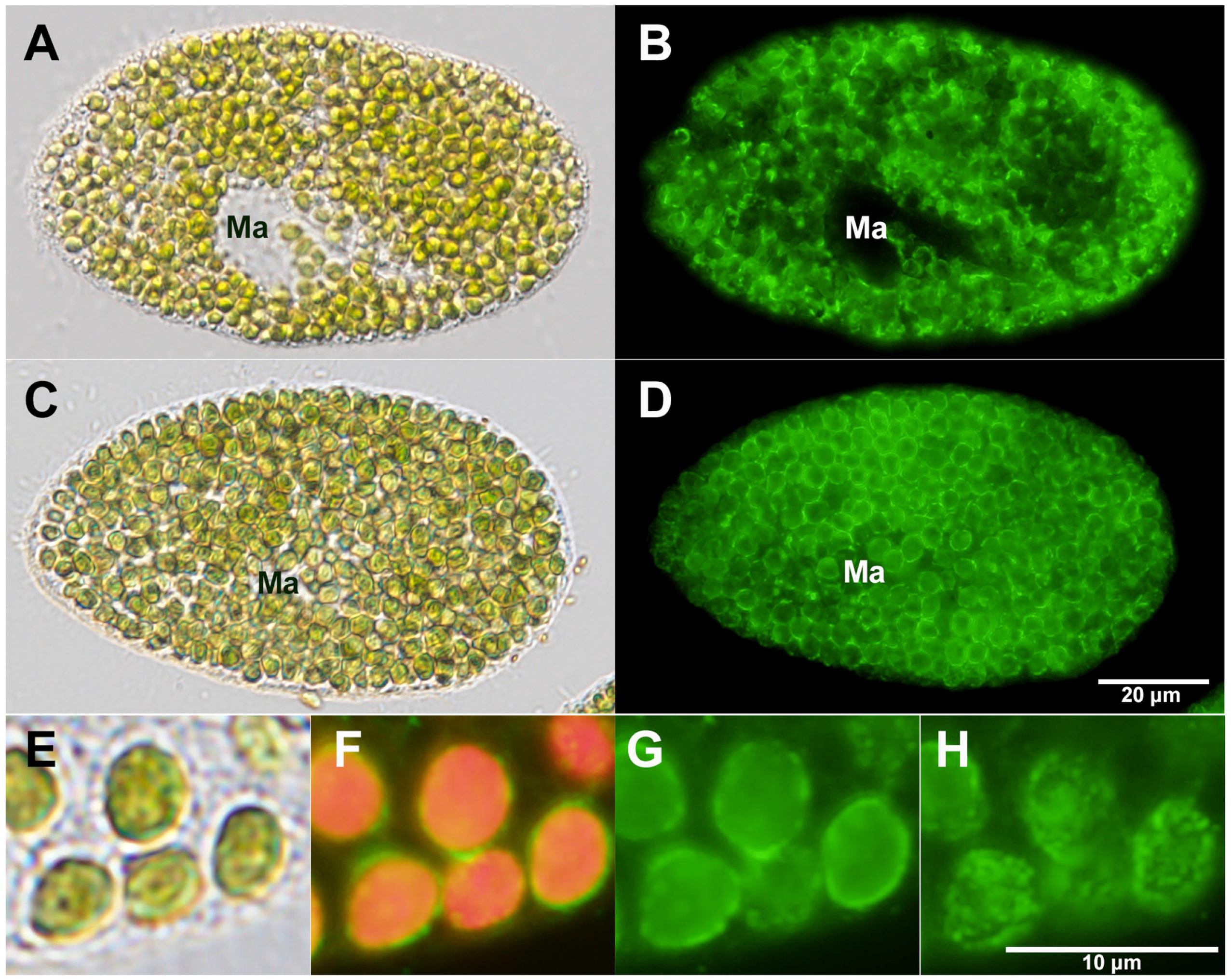
Effect of fixation methods on the visualization of the PV membrane antigen in *P. tritobursaria* Yad1g1N. (**A**, **C**) DIC images of cells fixed with 4% (v/v) PFA (**A**) or 99.5% (v/v) ethanol (**C**). (**B**, **D**) Corresponding indirect immunofluorescence images showing diffuse staining after PFA fixation (**B**) and sharply defined ring-like PV membrane signals after ethanol fixation (**D**). (**E**) High-magnification DIC image of symbiotic *Chlorella* sp. cells in ethanol-fixed Yad1g1N. (**F**) Merged image of PV membrane immunofluorescence (green) and algal autofluorescence (red). (**G**, **H**) Immunofluorescence images of the same field acquired at different focal planes. All images in panels E–H were obtained using a 100× objective lens. When the focal plane was adjusted to the algal surface (**H**), the PV membrane antigen signal appeared punctate, indicating that the apparently continuous signal observed at other focal planes may reflect the cumulative detection of discrete antigen-positive structures. Ma, macronucleus; Cy, cytopharynx.

### 3.2 Localization and retention of the PV membrane antigen

To characterize the antigen recognized by the anti-PV membrane mAb, we examined *P. tritobursaria* strains Yad1w and Yad1g1N (Figure 2). In the DIC image (Figure 2A), Yad1w cells were visible in the upper left corner, whereas the remaining cells corresponded to Yad1g1N. All samples were fixed using ethanol. Indirect immunofluorescence microscopy revealed strong fluorescence localized to the PV membrane surrounding endosymbiotic algae in Yad1g1N cells (Figure 2B), whereas no signal was detected in Yad1w cells, demonstrating that the antigen is specifically present in host cells harboring symbiotic algae. To determine whether the antigen persisted after algal isolation, *Chlorella* sp. cells isolated from Yad1g1N cells were examined (Figure 2C). The isolated algae displayed strong red autofluorescence (Figure 2D), and no fluorescence was detected around the algal surface using indirect immunofluorescence microscopy (Figure 2E), except for slight nonspecific signals in vacuoles. These observations indicate that the antigen was not retained after algal isolation, suggesting that the PV membrane was detached or lost during the isolation process. High-magnification observations of intact Yad1g1N cells revealed that all endosymbiotic algae positioned beneath the host cell cortex (white arrowheads) were consistently surrounded by a strongly fluorescent PV membrane (Figures 2F–H). Enlarged images showed that healthy green algae were uniformly enclosed by antigen-positive membranes (Figure 2I–K), whereas digested brown algae (black arrowheads) exhibited no detectable antigen signal, suggesting that PV membrane identity depends on the physiological state of the enclosed algae. Cells partially disrupted during sample preparation provided further insights into antigen persistence. Strong fluorescence remained within the damaged regions of ruptured Yad1g1N cells (Figures 2L–N), despite the absence of algae (white arrow). Similarly, membrane fragments remaining after cell disruption were immunopositive (Figures 2O–Q). Even in severely ruptured cells, in which only fragments of the host cell cortex were preserved, the antigen remained detectable (Figures 2R–T). These observations indicate that the PV membrane antigen persists independently of the presence of algae and is stably associated with host-derived cortical membrane structures.

**FIGURE 2.**
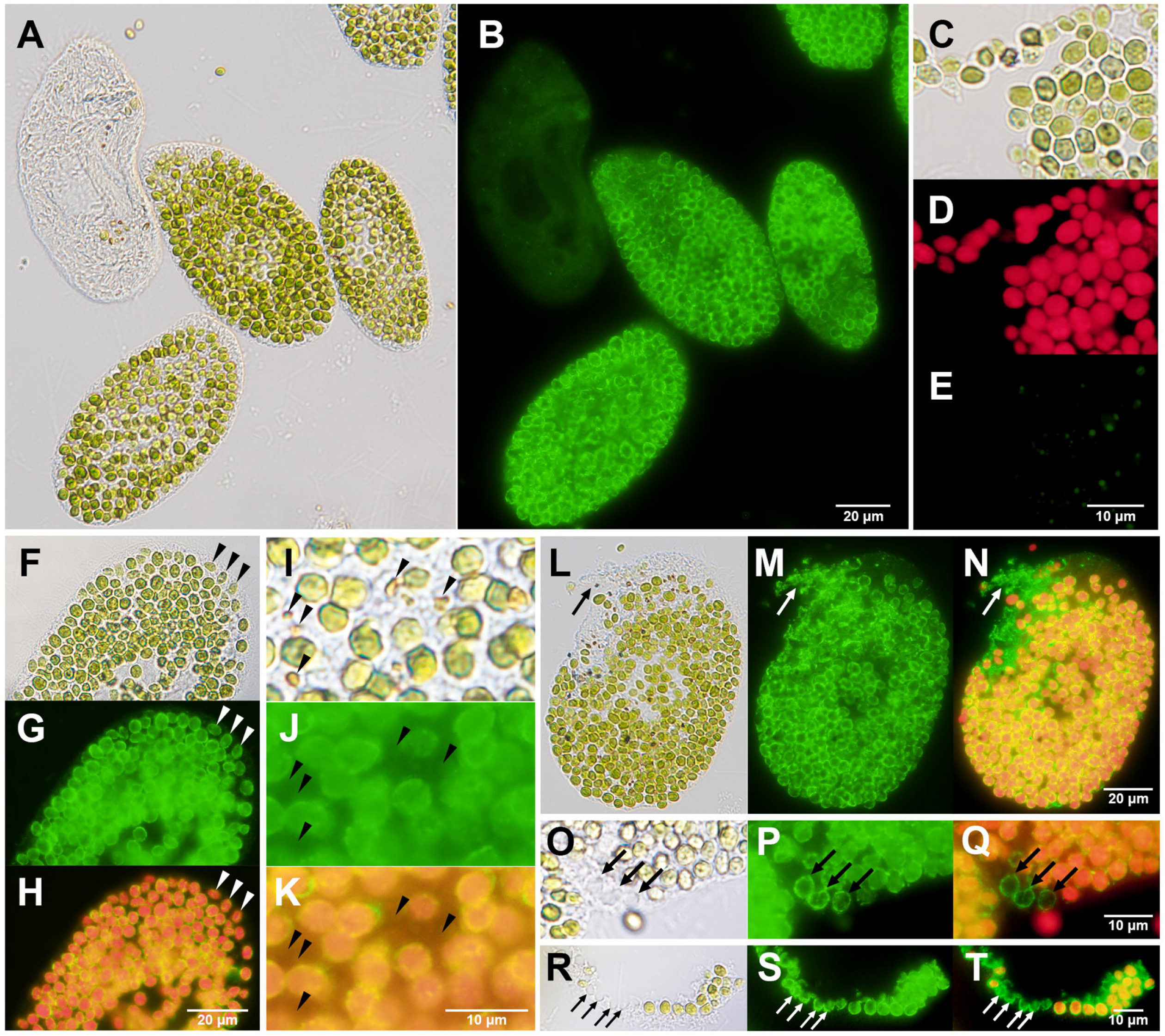
Immunofluorescence localization and retention of the PV membrane antigen in *P. tritobursaria*. (**A**, **B**) Mixed population of ethanol-fixed alga-free Yad1w (upper left) and algae-bearing Yad1g1N cells observed by DIC (**A**) and indirect immunofluorescence (**B**). Strong PV membrane signals were detected only in Yad1g1N. (**C**–**E**) Isolated symbiotic *Chlorella* sp. cells observed by DIC (**C**), autofluorescence (**D**), and immunofluorescence (**E**). No PV membrane signals were detected around the isolated algae. (**F**–**K**) Intact Yad1g1N cells showing PV membrane localization. Algae positioned beneath the host cortex were uniformly surrounded by antigen-positive PV membranes (**F**–**H**), whereas digested algae lacked detectable signals (**I**–**K**). (**L**–**T**) Disrupted Yad1g1N cells. Antigen-positive PV membrane structures persisted in partially ruptured cells (**L**–**N**), disrupted regions of the host cell lacking algae (**O**–**Q**), and severely disrupted cells (**R**–**T**), even in the absence of algae. Merged images show PV membrane immunofluorescence (green) and algal autofluorescence (red).

The same PV membrane antigen was also detected in another algae-bearing strain, YKK3g1N (Supplementary Figure 1), indicating that this antigen is not specific to Yad1g1N but is conserved across multiple *P. tritobursaria* strains.

### 3.3 Time-dependent emergence of the PV membrane following algal uptake

To quantitatively assess the timing of PV membrane maturation, we quantified the detection frequency of PV membrane antigen–positive membrane structures at each time point after algal uptake across five independent experiments (Table 1). PV membrane antigen–positive signals were not detected at early time points (10–30 min, 1.5–2.5 h, and 24 h), despite the formation of PV membranes by 24 h, indicating that newly formed PV membranes initially lack antigenicity. In contrast, antigen-positive PV membrane structures first appeared at 48 h and were detected surrounding all symbiotic algae by 72 h after infection.

**Table 1.** Detection frequency of PV membrane antigen–positive membrane structures at different stages after algal uptake.

| Time after mixing *Chlorella* sp. with alga-free *P. tritobursaria* | DV membrane, % (95 % CI), N | PV membrane, % (95 % CI), N |
| --- | --- | --- |
| 10–30 min | 0 (0–2.0) %, N=185 | Not observed at this time point |
| 1.5–2.5 h | 0 (0–1.8) %, N=205 | 0 (0–12.0) %, N=29<br>Rarely observed at this time point |
| 24 h | 0 (0–9.5) %, N=37<br>Rarely observed at this time point | 0 (0–0.1) %, N=2,481 |
| 48 h | 0 (0–23.2) %, N=14<br>Rarely observed at this time point | 20.0 (18.6–21.5) %, N=2,971 |
| 72 h | 0 (0–70.7) %, N=3<br>Rarely observed at this time point | 100 (99.9–100) %, N=4,347 |

The table summarizes the detection frequency of membrane structures showing positive immunofluorescence for the PV membrane antigen. DV membranes represent transient digestive vacuole-derived structures observed during early algal uptake, whereas PV membranes stably surround symbiotic algae and persist as antigen-positive structures. Exact 95% confidence intervals (CI) were calculated using the Clopper–Pearson exact method based on the binomial distribution. N indicates the number of membrane structures analyzed, not the number of host cells. “Not detected” or “rarely detected” indicates that PV membranes were not yet formed at early time points, whereas DV membrane structures became scarce at later stages. Data were obtained from five independent experiments.

Figure 3A summarizes the quantitative time course of PV membrane antigen acquisition after mixing alga-free *P. tritobursaria* with symbiotic *Chlorella* sp. The proportion of antigen-positive PV membranes remained at 0% until 24 h after mixing, increased at 48 h, and reached 100% at 72 h. The exact 95% confidence intervals were calculated using the Clopper–Pearson method. Throughout the observation period, the membranes of the DVs containing multiple algae remained consistently negative for the PV membrane antigen.

**FIGURE 3.**
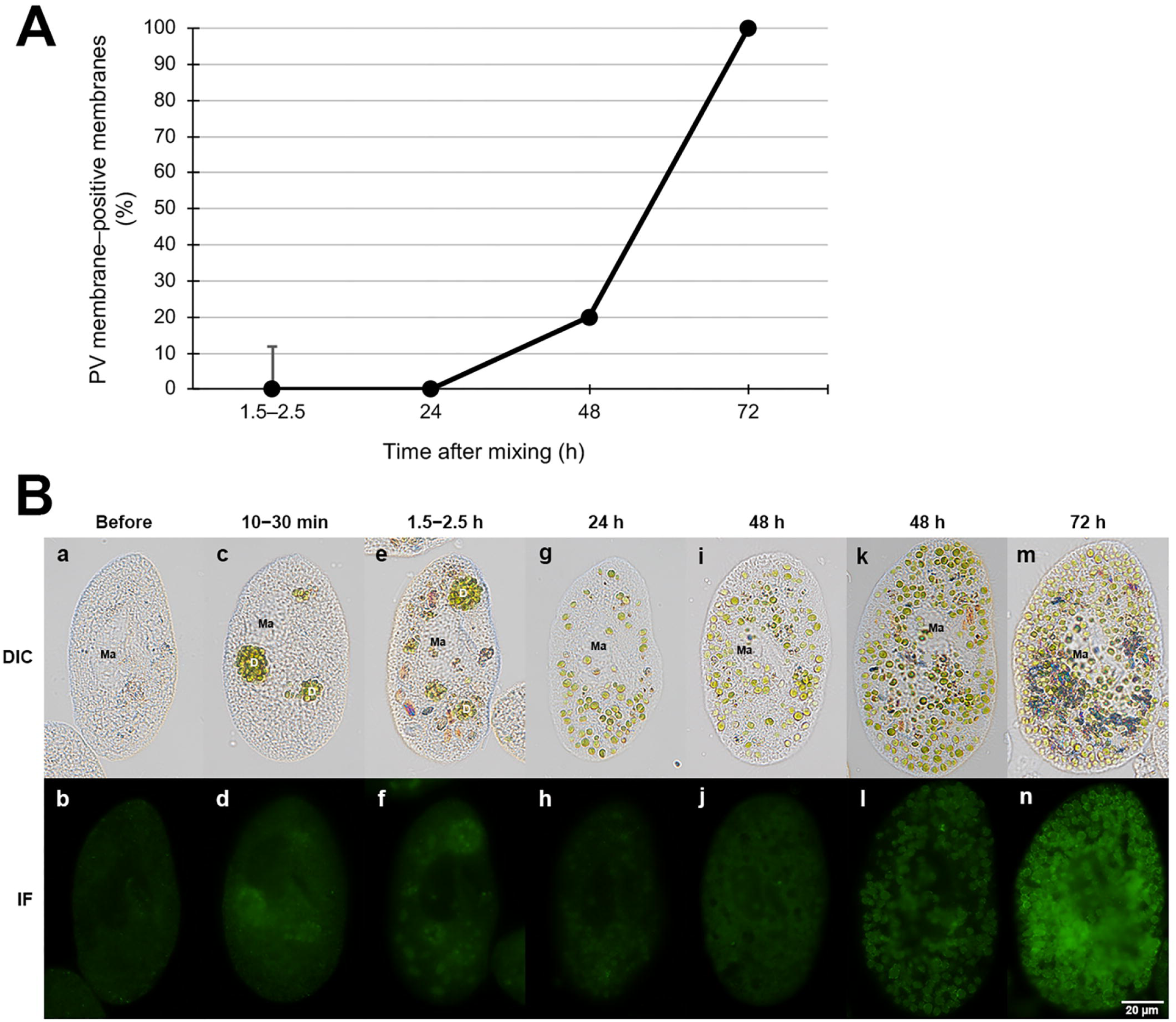
Time-dependent emergence of PV membranes in *P. tritobursaria* after mixing with *Chlorella* sp. (**A**) Proportion of PV membrane–positive membrane structures quantified by indirect immunofluorescence microscopy at the indicated time points after mixing. Error bars represent the 95% confidence intervals (CI) calculated using the Clopper–Pearson exact method. (**B**) Representative DIC and indirect immunofluorescence (IF) images showing time-dependent-changes in PV membrane detection after mixing. Alga-free *P. tritobursaria* Yad1w cells before mixing show no fluorescence signals (**a, b**). At 10–30 min (**c, d**) and 1.5–2.5 h (**e, f**), multiple algal cells were enclosed within DVs, as indicated by D in the DIC images. At 24 h (**g, h**), most internalized algae were individually positioned beneath the host cell cortex, but no PV membrane antigen signals were detected. At 48 h (**i–l**), PV membrane signals were detected around individually enclosed algae in a subset of host cells. At 72 h (**m, n**), PV membrane signals were detected around most individually enclosed algae. Green fluorescence indicates the PV membrane antigen detected by indirect immunofluorescence. Ma: macronucleus.

To visually corroborate the quantitative analysis, representative immunofluorescence images were obtained at each time point after mixing (Figure 3B). In alga-free Yad1w cells before mixing, no specific fluorescence signal was detected, confirming the absence of the PV membrane antigen in the alga-free state (Figures 3Ba, b). At 10–30 min and 1.5–2.5 h after mixing, large DVs containing multiple internalized algae were clearly observed using DIC microscopy (Figures 3Bc, e). As expected, no PV membrane antigen signals were detected around the ingested algae at early time points (Figures 3Bd, f), indicating that the DV membranes were antigen-negative. Twenty-four hours after mixing, most of the internalized algae were no longer enclosed within large DVs but were individually positioned beneath the host cell cortex, which is a characteristic feature of PV membranes. However, no specific PV membrane antigen signals were detected by immunofluorescence (Figures 3Bg, h). At 48 h after mixing, faint but discernible PV membrane signals became apparent in a subset of host cells, consistent with the emergence of PV membranes.

However, antigen-positive signals were not uniformly observed; most host cells remained largely antigen-negative (Figures 3Bi, j), whereas some cells exhibited detectable PV membrane fluorescence (Figures 3Bk, l). Clear discrimination of PV membrane antigen positivity is often difficult at this transitional stage, suggesting an asynchronous or heterogeneous onset of PV membrane maturation among host cells. At 72 h after mixing, strong PV membrane signals were stably observed surrounding nearly all individually enclosed algae within the host cells, indicating the establishment of uniform antigen-positive PV membranes (Figures 3Bm, n).

Under established short-term algal uptake conditions (1.5 min algal exposure), stable photoendosymbiosis was reproducibly achieved by 72 h, which we defined as the infection stage (Kodama and Fujishima, 2005). Importantly, throughout the early post-uptake process, prior to stable infection, DVs containing multiple algae remained consistently negative for PV membrane antigen. In contrast, acquisition of the PV membrane antigen occurs only after the transition from DV-type compartments to individually enclosed PV membranes. Notably, antigen-positive PV membranes did not exhibit a continuous or gradual increase over time; instead, they started to appear between 24 and 48 h after infection, indicating that PV membrane maturation proceeds through discrete transitions rather than a continuous process. Because all fluorescence images were acquired under identical exposure and staining conditions, the weak and heterogeneous signals observed at 48 h reflected genuine biological variability in PV membrane maturation, rather than technical detection limits.

Across three independent experiments, 288–528 host cells were examined per experiment, and 56.1–62.5% of the cells consistently harbored multiple green algae stably attached beneath the host cell cortex. These results confirm that stable endosymbiotic infection is reproducibly established by 72 h under short-term mixing conditions (1.5 min), as originally described by Kodama and Fujishima (2005), providing a reliable experimental framework for analyzing PV membrane maturation.

### 3.4 Induction of SPVS and its effect on the PV-membrane antigen

To examine the behavior of the PV membrane antigen during synchronous PV swelling (SPVS), Yad1g1N cells were treated with cycloheximide under continuous light. In untreated control cells (Figure 4A), the PV membrane closely adhered to the algal cell wall, and therefore, the PV membrane could not be visualized by DIC optics. In contrast, treatment with 10 µg/mL cycloheximide for 2 days induced SPVS, resulting in a clear separation between the PV membrane and the algal cell wall (Figure 4B, arrows). This separation created a visible gap that enabled the direct observation of the PV membrane in living cells, as previously described by (Kodama and Fujishima, 2008a). Numerous brown, degraded algae, indicative of digestion, were also observed at this stage. Previous TEM analyses have demonstrated that even after SPVS induction, each symbiotic alga remains individually enclosed by a PV membrane without fusion into a common vacuolar compartment (Kodama et al., 2011). To evaluate the behavior of the PV membrane antigen under the same conditions, cells were fixed and processed for indirect immunofluorescence microscopy (Figure 4C–F). In the control cells, strong fluorescence signals were detected around all symbiotic algae (Figure 4E). Although the PV membrane itself was not visible in the DIC images of fixed cells after SPVS induction (Figure 4D), indirect immunofluorescence signals were still detected surrounding the green algae (Figure 4F). After SPVS induction, PV membrane antigen signals often appeared as punctate structures along the swollen PV membranes. This punctate distribution is clearly illustrated in the enlarged inset (Figure 4F, upper right), which highlights multiple discrete fluorescent signals localized to the PV membrane surrounding individual algae. In contrast, brown digested algae (black arrowheads in Figure 4D) corresponded to regions lacking fluorescence signals (white arrowheads in Figure 4F), indicating that the PV membrane antigen is absent from membranes surrounding digested algae. Together, these results demonstrate that the PV membrane antigen remains associated with the PV membrane after SPVS induction as long as the enclosed algae remain intact but is lost from membranes surrounding algae undergoing digestion.

**FIGURE 4.**
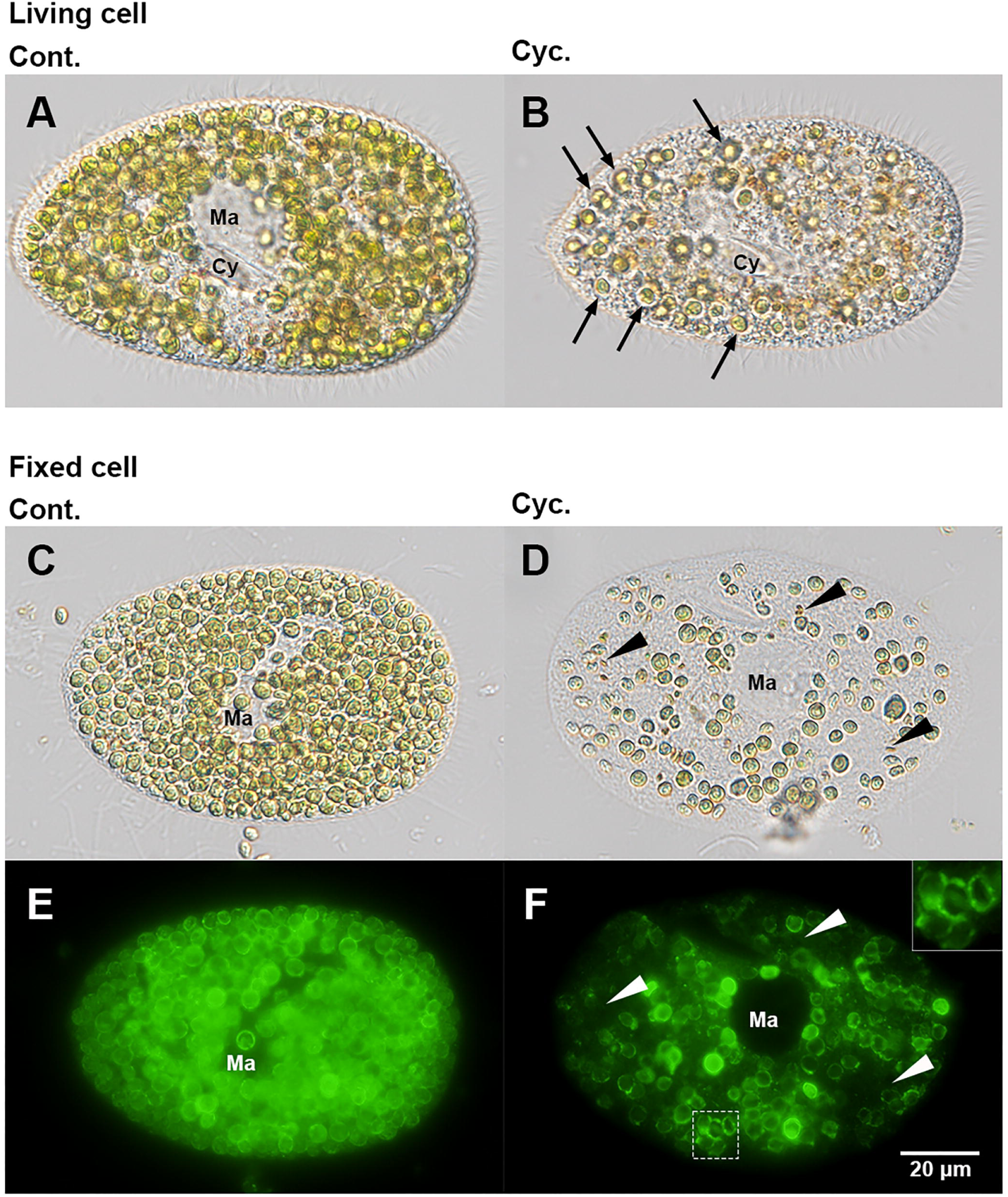
Induction of synchronous SPVS and localization of the PV membrane antigen in *P. tritobursaria* Yad1g1N. (**A**) DIC image of a control living cell. The PV membrane is not visible due to its close association with the algal cell wall. (**B**) DIC image of a living cell treated with cycloheximide (Cyc.). Arrows indicate regions exhibiting SPVS, where the separation between the PV membrane and algal cell wall is evident. Brown structures represent digested algae. (**C, D**) DIC images of control (**C**) and SPVS-induced (**D**) cells after fixation. Black arrowheads in (**D**) indicate digested algae. (**E**, **F**) Corresponding indirect immunofluorescence images of (**C**) and (**D**). Strong fluorescence signals are observed around intact algae under both conditions but are absent around digested algae after SPVS induction (white arrowheads). In SPVS-induced cells, fluorescence signals often appear as punctate structures along the PV membrane. The inset in (**F**) (upper right) shows a magnified view of the boxed region, highlighting the punctate localization of the PV membrane antigen. Ma, macronucleus; Cy, cytopharynx.

A quantitative analysis of PV membrane antigen retention before and after SPVS induction was performed using data obtained from three independent experiments, and the results are summarized in Table 2. Prior to SPVS induction, all PV membranes enclosing green symbiotic algae were positive for PV membrane antigen (100%, 95% CI: 99.84–100%). Antigen positivity was fully retained in PV membranes enclosing green or pale-green algae after SPVS induction (100%, 95% CI: 99.62–100% and 98.08–100%, respectively). In contrast, PV membranes surrounding digested brown algae after SPVS induction were consistently antigen-negative (0%, 95% CI: 0–2.58%). These results show that the PV membrane antigen is retained on PV membranes enclosing green or pale-green algae during SPVS, whereas it is absent from membranes surrounding digested brown algae.

**Table 2.**
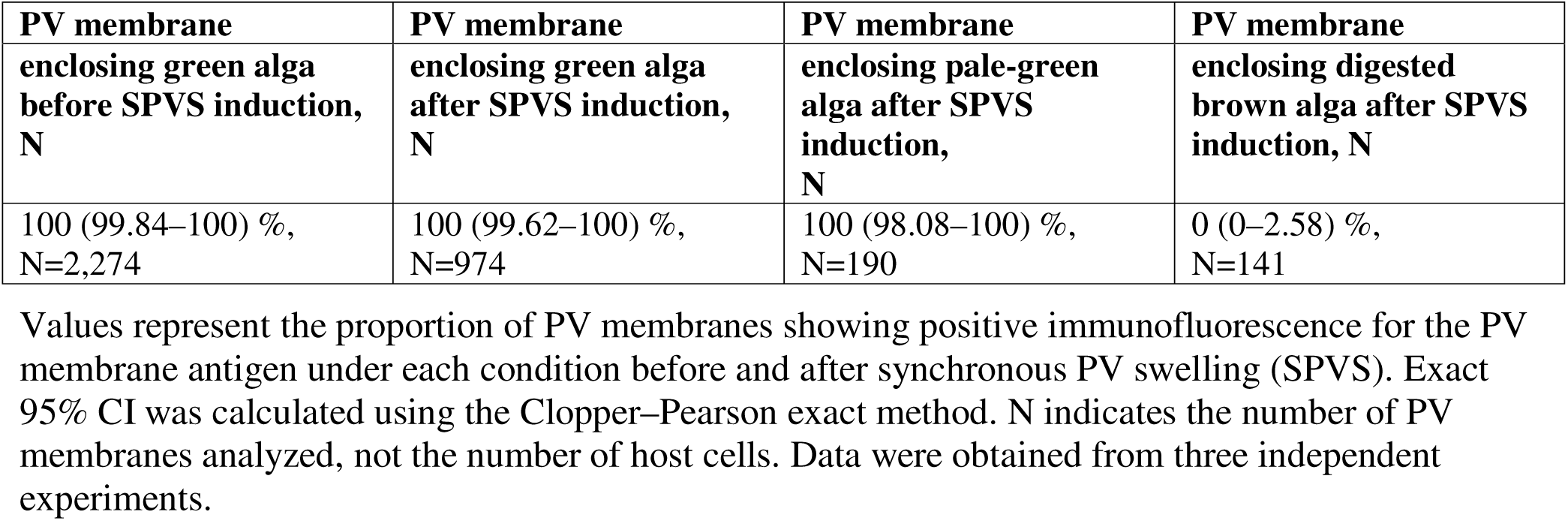
Proportion of PV membranes positive for the PV membrane antigen before and after synchronous PV swelling (SPVS).

## 4 Discussion

Our results show that in *P. tritobursaria*, the acquisition of a mature PV membrane occurs as a temporally regulated event after algal uptake, rather than as an immediate consequence of internalization. In this study, we combined membrane-level quantification with immunofluorescence imaging using a PV membrane-specific mAb to examine the temporal dynamics of DV and PV membrane formation following algal uptake. Throughout this study, the term “PV membrane antigen” refers to an antigen detected in close association with the PV membrane; however, the present data do not allow us to conclude that the antigen is uniformly distributed over the entire membrane surface. Quantitative analysis revealed a distinct transition window between 24 and 48 h after mixing, with complete acquisition of the PV membrane antigen within 72 h (Figure 3). In particular, our results provide the first direct molecular evidence addressing process (iii) of the previously proposed algal artificial re-photoendosymbiosis process, namely, maturation of the DV-derived membrane into a functionally and molecularly distinct PV membrane. These findings indicate that PV membrane formation represents a critical and temporally regulated checkpoint at which internalized *Chlorella* sp. escapes digestion and establishes a stable endosymbiotic state in *P. tritobursaria*. In this context, As previously reported (Kodama and Fujishima, 2005), DV membranes were observed only transiently during the early stages of algal infection. In contrast, once formed, PV membranes persist as stable and discernible structures, enabling a reliable temporal analysis of their appearance. The abrupt increase in PV membrane-positive structures between 24 and 48 h after algal uptake indicates that the DV-to-PV transition occurs within a defined temporal window and is actively regulated rather than simply resulting from the prolonged intracellular residence of the algae. Such stepwise and checkpoint-dependent membrane maturation has been well documented for phagosomes and pathogen-containing vacuoles in other eukaryotic systems, where discrete molecular transitions determine vacuolar fate rather than continuous progression (Flannagan et al., 2012; Levin et al., 2016).

In addition to temporal regulation, PV membrane antigens exhibit remarkable structural persistence within host cells. As shown in Figure 2, the antigen remained clearly detectable even in severely disrupted cells, in which symbiotic *Chlorella* sp. cells were absent, and only fragments of the host cortex were preserved. This observation indicates that the antigen is not directly associated with the algal surface or algal-derived components but localizes to a host-derived PV membrane structure that is stably linked to the cortical architecture of *P. tritobursaria*. Such localization is consistent with previous TEM studies, which have shown that mature PV membranes are positioned immediately beneath the host cortex and frequently associate with host organelles, including the mitochondria and endoplasmic reticulum (ER) (Song et al., 2017; Kodama and Fujishima, 2022).

Taken together, these findings support the view that the mature PV membrane functions as a host-organized symbiotic interface that is structurally integrated with the cortical and organellar architecture of the host cell, rather than as a transient membrane compartment defined solely by the presence of symbiotic algae. Consistent with this view, high magnification immunofluorescence imaging occasionally revealed punctate distributions of the PV membrane antigen along the PV membrane (Figure 1H), suggesting that the antigen may be organized in discrete domains or higher order membrane-associated assemblies rather than being uniformly distributed.

Although the mAb signal was detected in association with PV, the present observations do not allow us to conclude that the antigen is uniformly distributed over the entire PV membrane. Optical sectioning revealed punctate signals on the PV surface, suggesting that the antigen may be localized to discrete structures associated with the PV rather than the membrane as a whole. One possible interpretation is that the antigen is concentrated in specialized domains related to mitochondrial association with the PV, as suggested by previous ultrastructural observations. However, whether the antigen resides in the PV membrane itself or in discrete granular structures associated with mitochondrial attachment sites could not be resolved by the present study. Further analyses using immunoelectron microscopy or indirect immunofluorescence of PV-enclosed *Chlorella* sp. isolated by the Percoll-based method described by (Fujishima and Nishiyama, 2026), combined with mitochondrial markers, are required to clarify the precise sub-PV localization of the antigen.

A key technical and conceptual challenge in studying PV membranes has been the lack of suitable molecular tools. Because *P. bursaria* is a highly divergent non-model eukaryote and the PV membrane represents a symbiosis-specific compartment that does not correspond to classical endosomal or lysosomal membranes, commercially available antibodies against known membrane markers are largely inapplicable. In this context, the mAb described herein provides a rare and critical molecular entry point for the analysis of PV membrane maturation and functional state. Importantly, the antigen recognized by this antibody does not merely label a structural membrane component but instead reflects a specific, functionally maintained membrane state, highlighting the limitations of morphology-based definitions alone. The concept of a host-derived symbiotic membrane defined by molecular markers rather than morphology alone has also been established in cnidarian–dinoflagellate symbioses using monoclonal antibodies (Wakefiel and Kempf, 2001), supporting the broader relevance of antibody-based identification of mature symbiotic membranes.

Although the digested *Chlorella* sp. cells remained enclosed by membranes morphologically derived from the PV membrane after SPVS, as shown previously (Kodama et al., 2011), no antigenic signal was detected around these algae (Figures 4D, F). These membranes retained the ability to enclose a single algal cell, which is a characteristic feature of PVs, but exhibited properties resembling those of DVs, including lysosomal fusion and cytoplasmic streaming. This indicates that the antigen is lost when the essential PV membrane-specific properties are compromised, even if the membrane retains PV-like morphology. Thus, the antigen represents a molecular signature of PV membrane functionality rather than a simple structural component.

Previous studies have demonstrated that AcPase activity decreases rapidly after algal escape from the DV and before the algae become stably anchored beneath the host cell cortex (Kodama and Fujishima, 2009). Similar AcPase-based approaches have recently been applied to visualize digestive responses in cnidarians, revealing lysosomal and phagolysosomal AcPase activities during feeding and early algal interactions in corals (Sasamoto et al., 2025). However, our immunocytochemical analysis revealed that the loss of AcPase activity and subcortical localization were insufficient for antigen appearance. Even at 24 and 48 h, when AcPase activity was absent and algae were already positioned beneath the host cell surface, antigen signals were scarce (Figure 3). These observations indicate that antigen expression is not linked to DV-to-PV conversion, early PV differentiation, or cortical attachment but instead requires additional maturation steps that occur later in the infection process. This interpretation is further supported by the following observations. Antigen appearance occurred reproducibly approximately 48–72 h after algal uptake, a timing that corresponds to the onset of several independently reported features of functional symbiosis. Previous studies have shown that the synchronization of circadian rhythms between *P. tritobursaria* and symbiotic *Chlorella* sp. begins approximately 2–3 days after algal reinfection (Miwa, 2009), after which the stress resistance of the host and other symbiont-derived functions emerge at 3–4 days (Kinoshita et al., 2009). In addition, our earlier cell wall labeling experiments demonstrated that all symbiotic algae completed at least one round of cell division within the PV membrane by day 3 (Kodama and Fujishima, 2012). The close temporal alignment of these phenomena suggests that antigen emergence marks the transition from simple intracellular persistence to a functionally integrated symbiotic state. The distinction between physical coexistence and functional integration has also been recognized in other mutualistic systems. In arbuscular mycorrhizal symbiosis, plants discriminate among fungal partners and stabilize cooperation through reciprocal resource exchange, indicating that stable mutualism is achieved only after functional contributions are established (Kiers et al., 2011). Taken together, these findings indicate that the antigen is neither an early infection marker nor a constitutive PV membrane antigen. Instead, it is best interpreted as a molecular indicator of a “mature PV membrane formed only after sustained physiological interaction between the host and symbiont is achieved. The delayed appearance of the antigen implies a dependency on the integration of algal metabolism, host nutritional status, and potentially reciprocal signaling between the two partners.

Accordingly, the PV membrane should be viewed not as a compartment formed immediately after algal escape from the DV, but as a membrane system that progressively acquires its functional identity during symbiosis establishment.

Notably, ethanol fixation preserved the PV membrane antigen more clearly than PFA fixation (Figure 1). As discussed in the general immunocytochemistry guidelines, PFA fixation induces extensive protein cross-linking that can mask certain epitopes, whereas alcohol-based fixation methods often better preserve membrane-associated or conformationally flexible epitopes by precipitating proteins without covalent cross-linking (see, for example, R&D Systems, “Appropriate Fixation of IHC/ICC Samples”). These properties are consistent with the idea that the antigen reflects a maturation-associated molecular state of the PV membrane, rather than a general structural membrane component.

To place our findings in a broader biological context, it would be informative to compare PV membrane maturation in *P. tritobursaria* with host-derived symbiosome membranes described in other symbiotic systems (Roth et al., 1988). A broader perspective of cnidarian systems suggests that host-derived symbiosome membranes serve as dynamic regulatory interfaces. In corals and sea anemones (anthozoans), the symbiosome membrane is enriched in V-type H-ATPase (VHA), which acidifies the lumen to approximately pH 4 and promotes a carbon-concentrating mechanism for algal photosynthesis. VHA inhibition reduces luminal acidity and photosynthetic O production, demonstrating host control over symbiont physiology (Barott et al., 2015; Barott et al., 2022).

Additionally, Rhesus channels (ayRhp1) facilitate NH and CO diffusion and exhibit diurnal regulation, with higher abundance during the day and lower abundance at night, consistent with daytime nitrogen delivery and nighttime nitrogen limitation to optimize symbiont metabolism (Thies et al., 2022). Previous proteomic and genetic studies have shown that symbiosome membranes recruit host-derived lysosomal components, such as transporters and membrane-associated proteins, indicating that host membrane remodeling is essential for functional symbiosis (Peng et al., 2010). In green *Hydra*, classic physiological and cell biological studies have proposed that transient shifts in the pH of the PV regulate algal growth and maltose release, whereby acidification suppresses algal proliferation and favors sugar export, whereas alkalinization promotes algal growth (Douglas and Smith, 1984; Rands et al., 1992).These comparative insights align with our finding that the PV membrane in *P. tritobursaria* acquires the antigen only after a temporal delay, implying a maturation process analogous to transporter recruitment and luminal conditioning documented in anthozoans and proposed for *Hydra*. Together, these parallels suggest that dynamic remodeling of host-derived symbiotic membranes may represent a conserved strategy across distant eukaryotic lineages. Notably, the delayed acquisition of PV membrane antigenicity observed in *P. tritobursaria* highlights a level of temporal regulation that has been difficult to resolve in multicellular symbioses. Therefore, this unicellular system provides unique experimental access to the stepwise maturation of host-derived symbiotic membranes. The absence of antigen from membranes surrounding digested algae after SPVS further indicates that PV membrane identity is not fixed but instead requires active maintenance. This phenomenon parallels symbiosis breakdown processes described in cnidarians during bleaching or dysbiosis, where symbiosome membranes revert toward a phagolysosomal identity as host cells re-engage in digestive pathways (Weis, 2008; Davy et al., 2012). Thus, the presence of antigen reflects an actively sustained PV membrane state that is dismantled once symbiosis collapses.

As summarized in Figure 5, we propose a stepwise model in which host-derived membranes enclosing internalized algae undergo regulated maturation. Immature or pre-PV membranes initially lack antigenicity and subsequently acquire both functional competence and antigen expression, culminating in the formation of a mature PV membrane by approximately 72 h after algal uptake.

**FIGURE 5.**
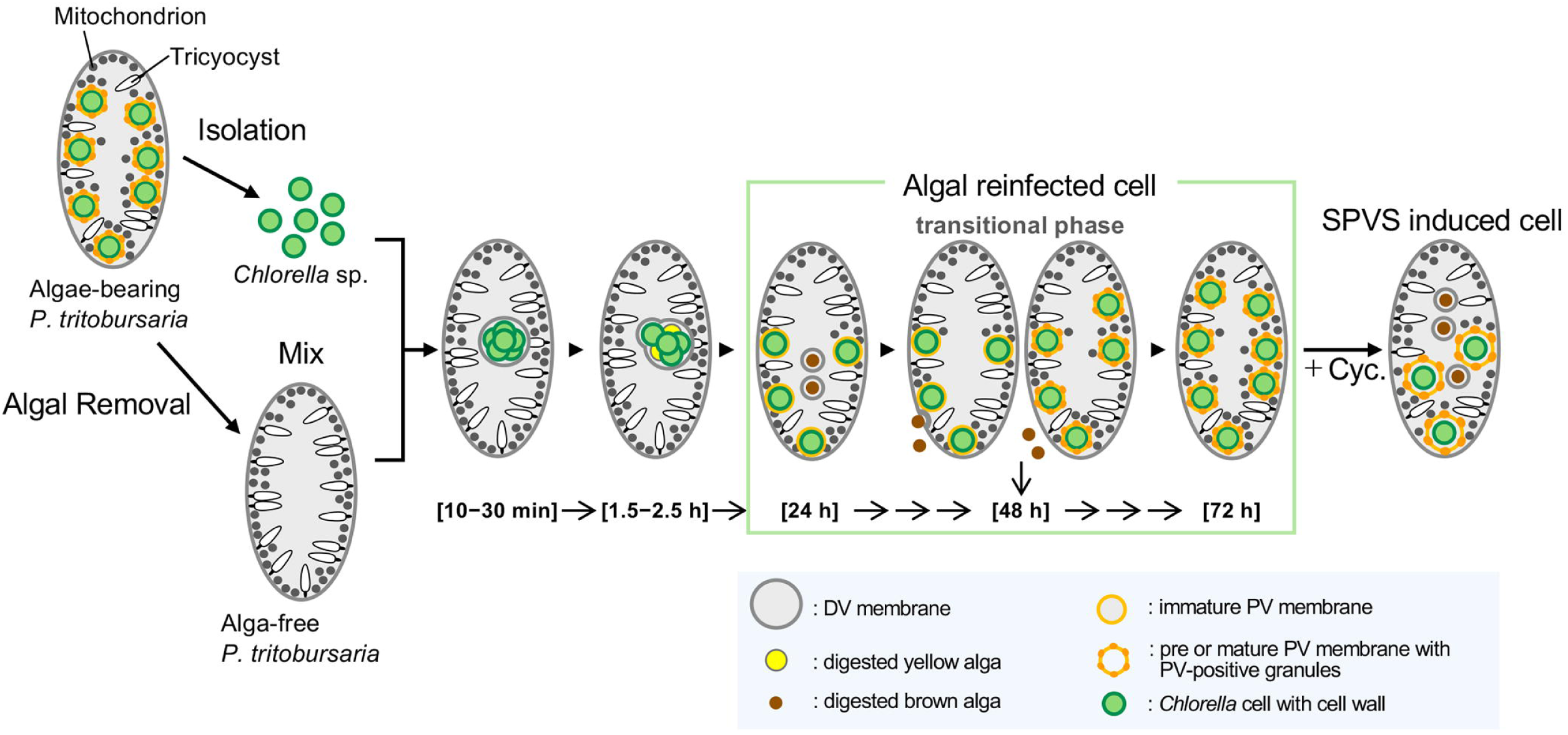
Conceptual model of stepwise PV membrane maturation and functional state transitions in *P. tritobursaria*. *Chlorella* sp. cells isolated from PV antigen–positive symbiotic *P. tritobursaria* were mixed with alga-free (aposymbiotic) *P. tritobursaria* cells to initiate reinfection. After mixing, *Chlorella* sp. cells are internalized into DVs, depicted with grey membranes. At early time points after algal uptake (10–30 min), internalized algae are enclosed within PV antigen-negative DVs. During the early post-uptake phase (1.5–2.5 h), DV membrane budding occurs; however, the resulting DV-derived membranes remain PV antigen-negative, and a subset of internalized algae exhibits early signs of digestion (yellow). By 24 h after algal uptake, individual algae become enclosed by antigen-negative immature PV membranes, depicted as light orange lines, which are positioned beneath the host cell cortex. At this stage, digestion has progressed further in many internalized algae, which are depicted predominantly as brown degraded algae. Previous studies have shown that in algae-bearing *P. tritobursaria* cells, trichocysts are locally reduced in regions occupied by symbiotic algae, whereas mitochondria tend to be distributed around the PV; however, the abundance of both organelles is reduced compared with alga-free cells (Kodama and Fujishima, 2011; 2022; Kodama and Fujishima, 2024). Accordingly, mitochondria and trichocysts are depicted near, but not in direct contact with, the PV membrane in this conceptual model, as their direct physical association during reinfection has not been determined. During the transitional stage (approximately 48 h after algal uptake), antigen-positive PV membranes are detected in a subset of host cells, whereas other cells remain antigen-negative, indicating heterogeneity in PV membrane maturation. At this time point, digested algal remnants are expelled from the host cell through the cytoproct. Cell division of the infected alga begins from 48 h after mixing. By 48 h after algal uptake, the presence of the PV membrane antigen is uniformly observed around mature PV membranes, depicted by orange outlines with dark orange granules. In *P. tritobursaria* cells that have established intracellular symbiosis with *Chlorella* sp., structurally stable adhesion is formed between the PV membrane and host mitochondria (Kodama and Fujishima, 2022). By 72 h after algal uptake, the presence of the PV membrane antigen is uniformly observed around mature PV membranes. Cycloheximide (Cyc.)-induced SPVS results in the retention of antigen positivity in the membranes surrounding intact or pale-green algae, whereas membranes enclosing digested brown algae lack detectable antigen. For clarity, *Chlorella* sp., mitochondria, and trichocysts are shown in reduced numbers to emphasize the temporal dynamics of PV membrane antigen acquisition rather than their actual abundance in host cells.

During this process, some ingested algae are degraded, consistent with previous observations (Kodama and Fujishima, 2005), indicating that the establishment of stable symbiosis involves selective retention rather than uniform survival of ingested algae. Importantly, antigen presence does not always coincide with a fully mature PV functional state; PV membranes surrounding green algae after SPVS can retain the antigen despite reverting to a DV-like or pre-PV identity, as shown in Fig. 4. Previous ultrastructural analyses have shown that symbiotic *Chlorella* sp. cells can appear green at the light microscopic level even when marked degeneration of chloroplast and nuclear structures is already evident by TEM, well before the onset of SPVS (Kodama et al., 2011). Taken together, our results reposition the PV membrane as a developmentally dynamic membrane, the functional maturation and molecular composition of which can be partially uncoupled from each other. The mAb described herein provides a powerful molecular marker for probing PV membrane maturation and offers new opportunities to dissect the mechanisms underlying the establishment, maintenance, and breakdown of symbiosis in protists.

## 5 Conclusion

In conclusion, this study demonstrates that PV membrane formation in *P. tritobursaria* is a temporally regulated process that constitutes a critical checkpoint during the establishment of photoendosymbiosis. Using a PV membrane-specific monoclonal antibody, we provided molecular evidence that PV membranes acquire their definitive symbiotic properties only after a defined post-uptake time window. This study establishes a framework for analyzing PV membrane differentiation at the molecular level. Although the molecular identity of the antigen remains to be determined, its specific and delayed appearance indicates that functional PV membrane formation is governed by a tightly controlled stepwise process. Identifying the antigen in future studies will further advance our understanding of PV membrane formation and maintenance and provide deeper insights into the cellular mechanisms underlying stable photoendosymbiosis.

## Conflict of Interest

The authors declare that the research was conducted in the absence of any commercial or financial relationships that could be construed as a potential conflict of interest.

## Author Contributions

Y.K.: Conceptualization, Data curation, Formal analysis, Funding acquisition, Investigation, Project administration, Resources, Supervision, Validation, Visualization, Writing – original draft; M.F.: Conceptualization, Funding acquisition, Investigation, Project administration, Resources, Supervision, Validation, Writing – review & editing.

## Funding

This study was supported by a Grant-in-Aid for Scientific Research (B) (grant number 23H02529) from the Japan Society for the Promotion of Science (JSPS) and the SDGs Research Project of Shimane University granted to Y.K., and by a Grant-in-Aid for Scientific Research (B) (grant number 26291073) from the Japan Society for the Promotion of Science (JSPS) and the Academist Crowdfunding in 2017 granted to M.F.

## Acknowledgments

We would like to express our sincere gratitude to Ryo Nabara of Department of Science and Engineering, Graduate School of Science and Engineering, Yamaguchi University, for his cooperation in culture of a hybridoma clone that produces monoclonal antibodies specific to PV membrane. This study was conducted as part of the SDGs Research Project at Shimane University, Japan. The authors used M365 Copilot (Microsoft; GPT-5-based model), Paperpal (version 4.29.1) and ChatGPT (OpenAI, GPT-5.5) to assist with English language editing, but all scientific contents and interpretations were determined by the authors. The authors take full responsibility for the content of the manuscript.

## Notes

### Competing Interest Statement

The authors have declared no competing interest.

