## Supplementary Figure 1 for "First molecular evidence that perialgal vacuole membrane maturation is temporally regulated during establishment of *Chlorella variabilis* photoendosymbiosis in *Paramecium tritobursaria*"

### *Supplementary Material*

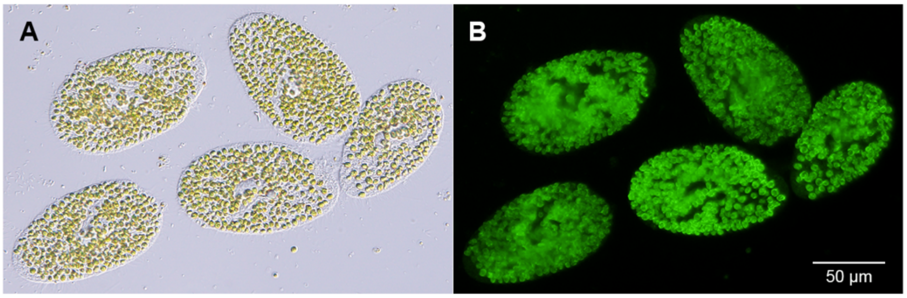


**Supplementary Figure 1.** Presence of PV membrane antigen in another algae-bearing strain (YKK3g1N; *P. tritobursaria*, mating type III). Ethanol-fixed YKK3g1N cells (**A**) showed strong AF488 fluorescence around all endosymbiotic *Chlorella* spp. (**B**), indicating that the antigen recognized by mAb is conserved across multiple *P. tritobursaria* strains. This finding demonstrates that the antigen is not specific to the Yad1g1N strain and suggests that the corresponding PV membrane molecule may be preserved among algae-bearing *P. bursaria* species.
